# The Epidemiology and Transmissibility of Zika Virus in Girardot and San Andres Island, Colombia

**DOI:** 10.1101/049957

**Authors:** Diana Patricia Rojas, Natalie E. Dean, Yang Yang, Eben Kenah, Juliana Quintero, Simon Tomasi, Erika L. Ramirez, Yendy Kelly, Carolina Castro, Gabriel Carrasquilla, M. Elizabeth Halloran, Ira M. Longini

## Abstract

**Background:** Zika virus (ZIKV) is an arbovirus in the same genus as dengue virus and yellow fever virus. ZIKV transmission was first detected in Colombia in September 2015. The virus has spread rapidly across the country in areas infested with the vector Aedes aegypti. As of March 2016, Colombia has reported over 50,000 cases of Zika virus disease (ZVD).

**Methods:** We analyzed surveillance data of ZVD cases reported to the local health authorities of San Andres, Colombia, and Girardot, Colombia, between September 2015 and January 2016. Standardized case definitions used in both areas were determined by the Ministry of Health and Colombian National Institute of Health at the beginning of the ZIKV epidemic. ZVD was laboratory-confirmed by a finding of Zika virus RNA in the serum of acute cases. We report epidemiological summaries of the two outbreaks. We also use daily incidence data to estimate the basic reproductive number R_0_ in each population.

**Findings:** We identified 928 and 1,936 laboratory or clinically confirmed cases in San Andres and Girardot, respectively. The overall attack rate for reported ZVD detected by healthcare local surveillance was 12·13 cases per 1,000 residents of San Andres and 18·43 cases per 1,000 residents of Girardot. Attack rates were significantly higher in females in both municipalities. Cases occurred in all age groups but the most affected group was 20 to 49 year olds. The estimated R_0_ for the Zika outbreak in San Andres was 1·41 (95% CI 1·15 to 1·74), and in Girardot was 4·61 (95% CI 4·11 to 5·16).

**Interpretation:** Transmission of ZIKV is ongoing and spreading throughout the Americas rapidly. The observed rapid spread is supported by the relatively high basic reproductive numbers calculated from these two outbreaks in Colombia.

**Funding:** This work was supported by National Institutes of Health (NIH) U54 GM111274, NIH R37 AI032042 and the Colombian Department of Science and Technology (Fulbright-Colciencias scholarship to D.P.R). The funding source had no role in the preparation of this manuscript or in the decision to publish this study.

**Research in Context:** *Evidence before this study:* The ongoing outbreak of Zika virus disease in the Americas is the largest ever recorded. Since its first detection in April 2015 in Brazil, around 500,000 cases have been estimated, and the virus is spreading rapidly in the Americas region. There are many unanswered questions about the transmissibility and pathogenicity of the virus. Limited data are available from recent outbreaks occurring in islands in the Pacific, and little epidemiological data is available on the current outbreak. We searched PubMed on March 12, 2016, for epidemiological reports on Zika virus outbreaks using the search terms “Zika” AND “Basic reproductive number”. We applied no date or language restrictions. Our search identified one previous paper assessing the basic reproductive number, *R*_0_ of Zika virus in Yap Island, Federal State of Micronesia and in French Polynesia, but no papers estimating *R*_0_ using data from the Latin American Zika outbreak. Because of the sparsity of the data, we could not do a detailed systematic review at this point in time.

*Added value of this study:* We report detailed epidemiological data on outbreaks in San Andres and Girardot, Colombia. Because such reports are currently unavailable, we provide early information on age and gender effects and the functioning of local and national surveillance in the second-most affected country in this epidemic. We provide early estimates of *R*_0_. Our results can be used by mathematical modelers to understand the future impact of the disease and potential spread.

*Implications of all the available evidence:* We report attack rates similar to those reported in the Yap Island outbreak. We find that Zika impacts individuals of all ages, though the most affected age group is 20 to 49 years of age. The surveillance system detected more cases among women in both areas, though this finding may be attributable to reporting bias. Our estimates of *R*_0_ imply that Zika has the capacity for widespread transmission in areas with the vector.

## 1. Introduction

First isolated in the Zika Forest of Uganda in 1947, Zika virus (ZIKV) is an arbovirus primarily transmitted by *Aedes aegypti* mosquitoes [1]. Although ZIKV has circulated in Africa and Asia since the 1950s, little is known about its transmission dynamics [2]. Recent outbreaks in Yap Island in the Federated States of Micronesia (2007), French Polynesia (2013), and other Pacific islands, including Cook Islands, Easter Island, and New Caledonia (2014), indicate that ZIKV has spread beyond its former geographic range [3, 4, 5, 6]. In April 2015 ZIKV was isolated in the Northeast of Brazil [7]. As of March 11, 2016, around 500,000 Zika virus disease (ZVD) cases have been estimated in Brazil, and autochthonous circulation has been observed in 31 countries in the Americas region. Further spread to countries within the geographical range of *Ae. aegypti* mosquitoes is considered likely [8].

ZIKV is a flavivirus in the same genus as dengue virus and yellow fever virus. Infection typically causes a self-limited dengue-like illness characterized by exanthema, low-grade fever, conjunctivitis, and arthralgia [9]. While illness is believed to be mild or asymptomatic in approximately 80% of the infections [10], an increase in rates of Guillain-Barré syndrome (GBS) has been observed during ZIKV outbreaks [8, 11, 12]. Furthermore, in October 2015, the Brazilian Ministry of Health reported a dramatic increase in cases of microcephaly in Northeast Brazil where ZIKV had been circulating [13]. On the basis of the possible link between ZIKV, GBS and microcephaly, the World Health Organization declared a public health emergency on February 1, 2016 [14].

The virus was first detected in Colombia in mid-September 2015 in a municipality called Turbaco on the Caribbean coast. Turbaco is located approximately 20 minutes from Cartagena, a well known commercial and tourist hub (Figure 1). In October 2015, ZIKV spread through the central region of the country, appearing in areas with endemic dengue and ongoing circulation of chikungunya (CHIKV) since 2014. Through March 2016, Colombia has reported over 50,000 cases of ZVD, making it the second-most affected country in this outbreak, after Brazil. There were 2,090 laboratory-confirmed cases with the rest being suspected cases or confirmed by clinical findings [15]. Up to March 2016, 280 cases of neurological complications including Guillain-Barré syndrome (GBS) and three deaths possibly associated with ZVD have been reported in Colombia [16]. As of March 2016, there have been several suspected but no confirmed cases of ZIKV-associated microcephaly in Colombia [15].

In this paper we describe local ZIKV outbreaks in Colombia in Girardot and San Andres Island between September 2015 and January 2016 for which detailed epidemiological data are available. We conduct an investigation to define the epidemiological features of these outbreaks and to estimate corresponding transmission parameters.

## 2. Methods

### San Andres

San Andres is the largest island in a Colombian archipelago in the Caribbean sea located about 750 km north of mainland Colombia and 230 km east of Nicaragua (Figure 1). The island has an area of 27 km^2^, a population of 54,513 inhabitants across 13,652 households, and a population density of 2,932 habitants per km^2^ in 2010 [17, 18]. The average temperature is 27·3°C, and 80% of the total annual rainfall of 1,700 mm occurs during the heavy rainy season between October and December. The weather is humid subtropical with occasional tropical cyclones and hurricanes. The population in San Andres has two main ethnic groups: Afro-Colombians (17.5%) and Raizal (an ethnic group of mixed Afro-Caribbean and British descent) (39.2%) [18]. The most productive breeding sites of *Ae. aegypti* in San Andres are unprotected water containers located in the households. San Andres has experienced low dengue transmission since 1983. Since 1995, the frequency of dengue outbreaks increased to every two to five years with a mean annual incidence of 43·6 cases per 100,000 inhabitants between 1999 and 2010 [17]. In 2014, CHIKV began circulating in San Andres, reaching an annual incidence of 365·1 cases per 100,000 inhabitants [19].

**Figure 1:**
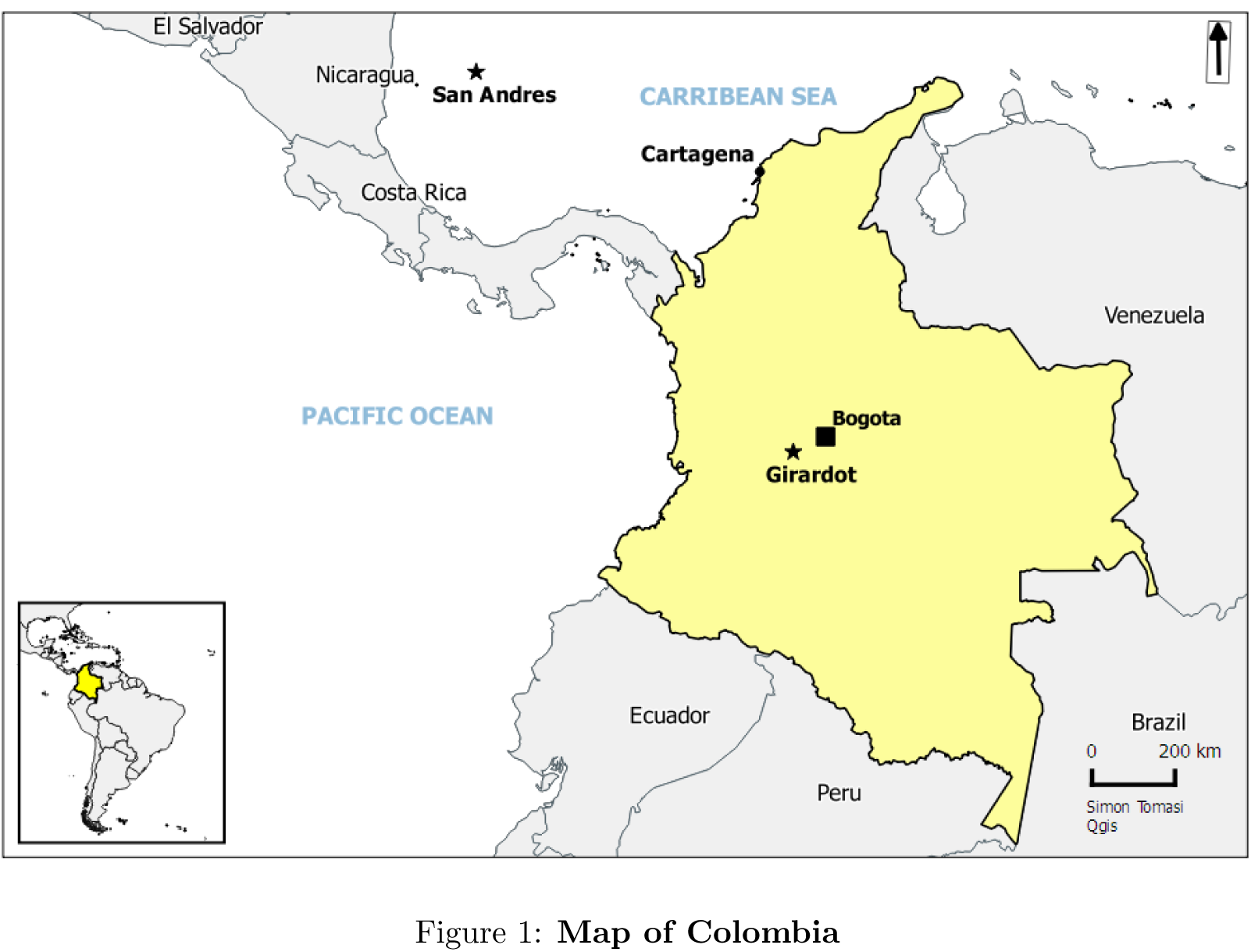
Map of Colombia

### Girardot

The city of Girardot is located 134 km (2 hours drive) from the capital city of Bogota (Figure 1). Girardot has 102,225 inhabitants across approximately 23,000 households based on the most recent census from the National Statistics Department (NSD) [20], though the population triples during weekends and holidays. Girardot is 289 meters above sea level. The average temperature is 33.3°sC, and the relative humidity is 66%. The mean annual precipitation is 1,220 mm with a rainy season extending from May through October [21]. The most productive breeding sites of *Ae. aegypti* in Girardot are unprotected private water containers, such as water storage tanks used in the households during the dry and rainy seasons, while public spaces provide more breeding sites during the rainy season [22]. Girardot has experienced hyperendemic transmission of dengue since 1990 with simultaneous circulation of all four serotypes; the mean annual incidence was 572·2 per 100,000 inhabitants between 1999 and 2010 [17]. In late 2014, CHIKV began circulating in Girardot reaching an annual incidence of 394 per 100,000 inhabitants in 2014 and 8,416 per 100,000 inhabitants in 2015.

### Case definition and laboratory analysis

We analyzed surveillance data from nine local health care sites in San Andres and twenty-two local health care sites in Girardot. Standardized case definitions used in both areas were defined by the Ministry of Health (MoH) and Colombian National Institute of Health (C-NIH) at the beginning of the ZIKV epidemic. A suspected ZVD case is defined as a person who lived or traveled in an area below 2,000 meters above sea level who presents with maculopapular exanthema, temperature higher than 37.2°C, and one or more of the following: non-purulent conjunctivitis, arthralgia, myalgia, headache, or malaise. A laboratory confirmed case is a suspected case with a ZIKV positive reverse-transcriptase-polymerase-chain-reaction (RT-PCR) result as determined by the C-NIH virology reference laboratory. A clinically confirmed case is a suspected case that lived or traveled in an area with laboratory confirmed ZIKV circulation prior to onset of symptoms [23].

At the start of the outbreak, all suspected cases were reported based on the suspected case definition to the Colombian national surveillance system. Once laboratory confirmation from C-NIH was performed for cases in Girardot and San Andres, the new suspected ZIKV cases were laboratory confirmed if they fell into the risk groups defined by the C-NIH: newborns, age <1 year, age >65 years, and cases with co-morbidities [23]. After ZIKV circulation was confirmed in the regions, all suspected cases were reclassified as clinically confirmed.

### Data collection

The data was collected using the C-NIH standard report form for Zika surveillance. The form includes socio-demographic variables. We analyzed a deidentified data set with the following variables: gender, age, pregnancy status, date of symptom onset, date the case visited the health care facility, date the case was reported to the national surveillance system, and case type (suspected, laboratory confirmed, clinically confirmed) [24].

### Statistical analysis

We calculated overall and age/gender-specific attack rates using population census data from NSD [20]. Surveillance data were analyzed using R version 3·2·0 [25]. For descriptive results, categorical variables are presented as proportions and continuous variables by the median and interquartile range (IQR) or range. The impact of age and gender on attack rates was tested using log-linear models for case counts with age category, gender, and an interaction as independent variables with population size as an offset.

To estimate the basic reproductive number *R*_0_ in each population, we used maximum likelihood methods to fit a chain-binomial model to daily incidence data [26]. *R*_0_ is the median effective reproductive number during the growth phase of the epidemic, after accounting for early under-reporting. (See Supplementary Online Materials for additional details on the model.)

## 3. Results

In San Andres, we identified 928 reported ZVD cases (Table 1). Of these cases, 52 (5·6%) were laboratory confirmed by RT-PCR on acute phase samples collected within five days of symptom onset, and 876 (94·4%) cases were clinically confirmed.

**Table 1:** Characteristics of reported cases of Zika Virus Disease

| Region | San Andres | Girardot |
| --- | --- | --- |
| Total number of cases | 928 | 1936 |
| Laboratory confirmed cases | 52 (5.6%) | 32 (1.7%) |
| Clinically confirmed cases | 876 (94.4%) | 1904 (98.3%) |
| Female | 589 (63.5%) | 1138 (58.8%) |
| Median age, years (IQR) | 31 (15, 47) | 34 (24, 46) |
| Median time, days, to visit healthcare facility from symptom onset (IQR) | 4 (1, 16) | 1 (1, 2) |

The dates of symptom onset among cases in San Andres ranged from September 6, 2015, to January 30, 2016 (Figure 2). Though the earliest case reported symptom onset on September, 6, 2015, the local health care authorities did not receive laboratory confirmation of ZIKV until October 22, 2015. The number of cases peaked in epidemiological week 45 (November 8 to 14) and subsided in the last week of December. The median time between symptom onset and visiting a health care facility was 4 days (IQR 1 to 16). 79% of cases were reported to the national surveillance system on the same day they visited the health care facility. The median age of reported ZVD cases in San Andres was 31 years old (IQR 15 to 47; range 12 days to 82 years). 589 (63·5%) of the reported cases occurred in females. During this time period 70 dengue cases and 10 CHIKV cases were confirmed in San Andres.

**Figure 2:**
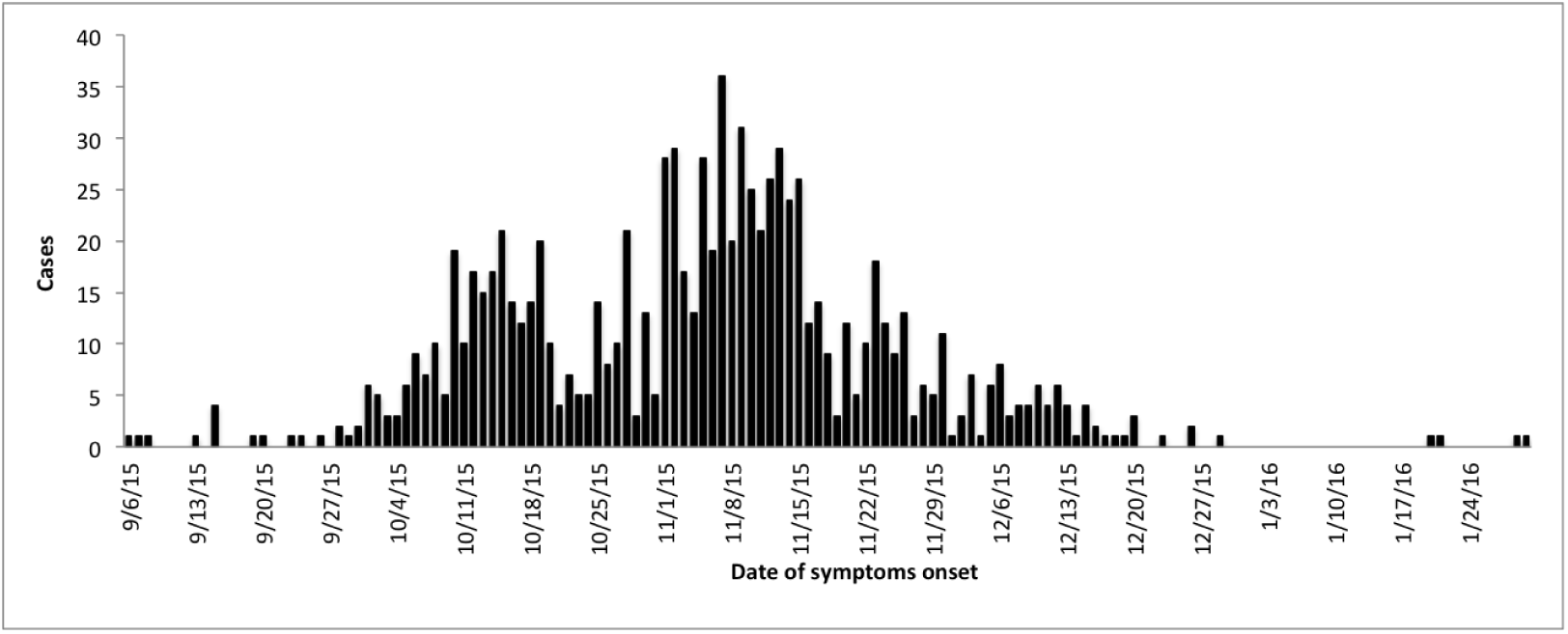
Daily ZVD incidence for San Andres, Colombia

The overall attack rate for ZVD reported by local surveillance was 12·13 per 1,000 San Andres residents. The gender-specific attack rates were 15·34 per 1,000 females and 8·91 per 1,000 males; the difference was significant adjusting for age (p<0·001). Cases occurred among all age groups, but the incidence of ZVD detected by local surveillance was highest among persons 20 to 49 years old (Figure 3); there was significant heterogeneity across the age groups (p<0·001). Attack rates were higher in females across all age groups 10 years old and above; there was a significant interaction between age and gender (p<0·001).

**Figure 3:**
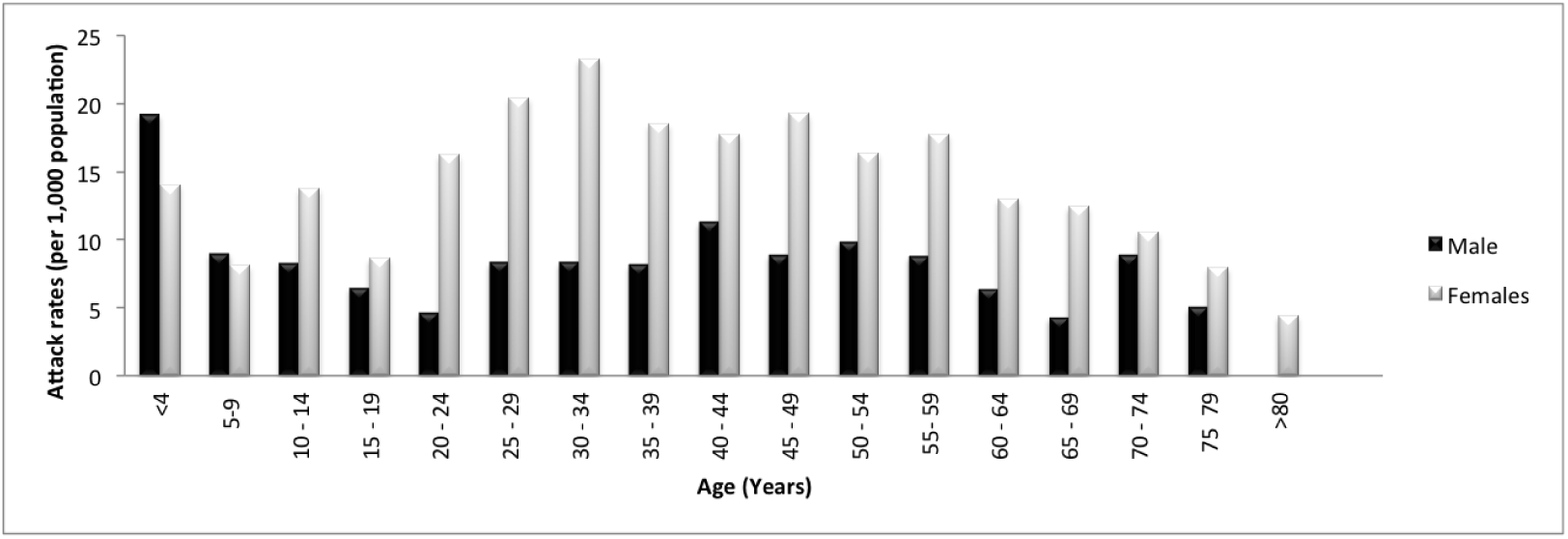
Age-and gender-specific ZVD attack rates for San Andres, Colombia

Thirty two pregnant women with ZVD were reported in San Andres and are being followed according to national guidelines. By March 2016, fourteen of them had given birth with no microcephaly reported. There were eight neurological syndromes reported in San Andres, including GBS and meningoencephalitis attributed to ZIKV and among them one death was reported. The incidence rate of neurological syndromes among ZVD cases in San Andres is 8·6 per 1,000 cases.

### Girardot

In Girardot, we identified 1,936 reported ZVD cases (Table 1). Of these cases, 32 (1·7%) were laboratory confirmed by RT-PCR on acute phase samples collected within five days of symptom onset and 1,904 (98·3%) were clinically confirmed. During this time period were confirmed 67 dengue cases and 37 CHIKV cases in Girardot.

The date of symptom onset among cases in Girardot ranged from October 19, 2015, to January 22, 2016 (Figure 4). The first suspected case was reported on October 23, 2015, with laboratory confirmation obtained on January 27, 2016. The number of cases peaked in epidemiological week 48 (November 29 to December 5) and subsided in early January. The median time between symptom onset and visiting a health care facility was 1 day (IQR 1 to 2 days). 89% of cases were reported to the national surveillance system on the same day they visited the health care facility. The median age of confirmed ZVD cases was 34 years old (IQR 24 to 46; range 15 days to 92 years). 1138 (58.8%) of the cases occurred in females.

**Figure 4:**
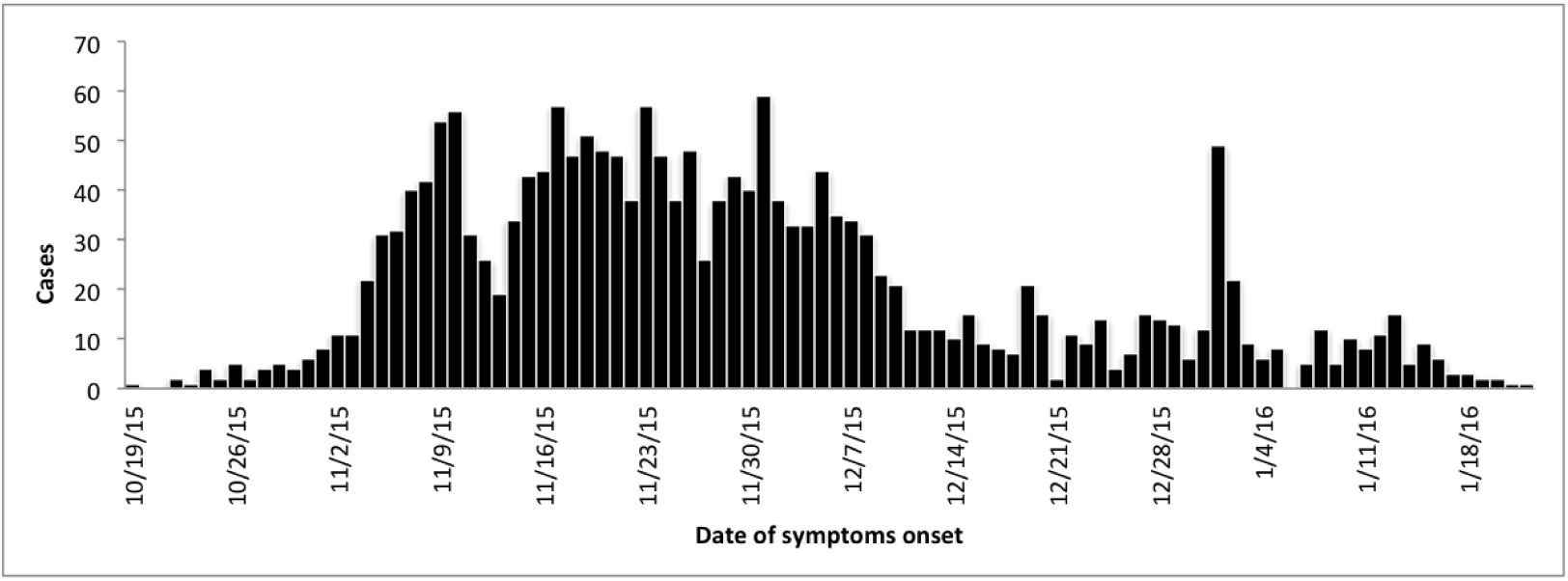
Daily ZVD incidence for Girardot, Colombia

The overall attack rate for confirmed ZVD detected by local surveillance was 18·43 per 1,000 Girardot residents. The gender-specific attack rates were 20·53 per 1,000 females and 16·07 per 1,000 males; the difference was significant adjusting for age (p<0·001). Cases occurred among all age groups, but the incidence of ZVD detected by local surveillance was highest among persons 20 to 49 years old (Figure 5); there was significant heterogeneity across the age groups (p<0·001). Attack rates were higher in females in all age groups except in those 10 to 14 and 65 to 69 years old; there was no significant interaction between age and gender.

**Figure 5:**
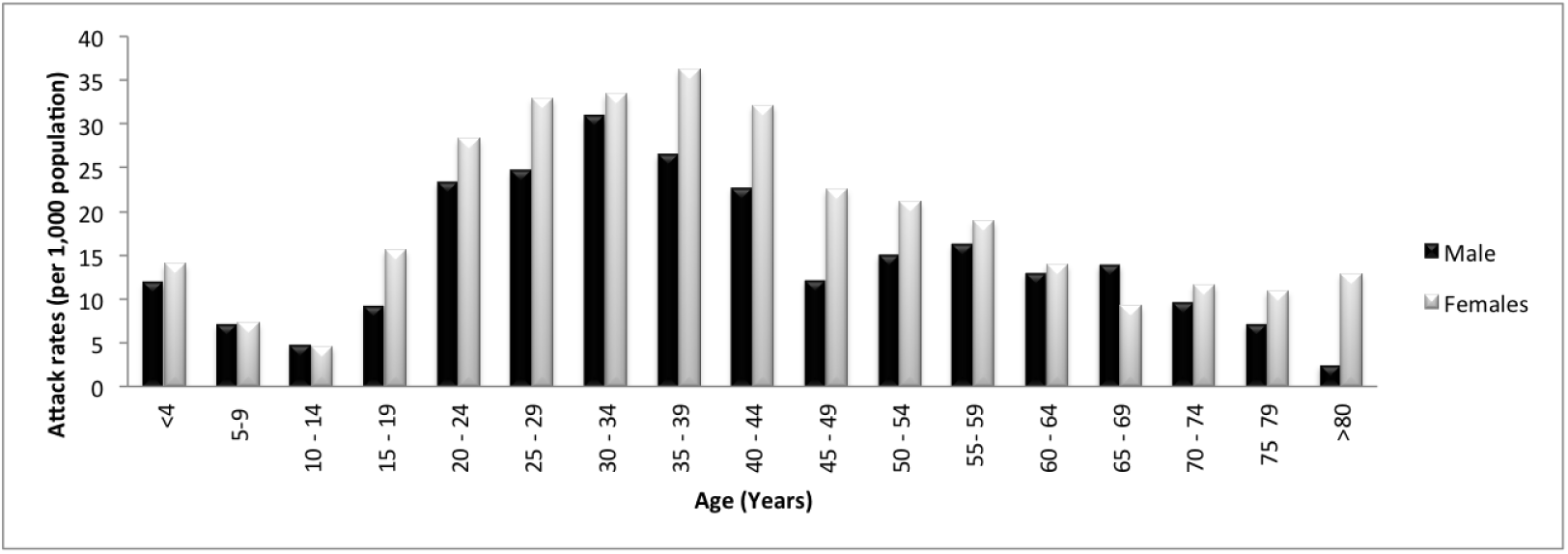
Age-and gender-specific ZVD attack rates for Girardot, Colombia

Sixteen pregnant women with ZVD were reported in Girardot and are being followed according to national guidelines. By March 2016, seven of them had given birth with no complications or microcephaly reported. Nine cases with GBS have been reported after an initial suspected ZIKV infection; laboratory-confirmation of ZIKV is pending. There were no deaths attributed to ZIKV. The incidence rate of neurological syndromes among ZVD cases in Girardot is 4·6 per 1,000 cases.

### Basic reproductive number calculations

The basic reproductive number (*R*_0_) was estimated using daily incidence data. The model assumes a mean serial interval (time between successive cases in a chain of transmission) of 22 days, based on a mean incubation period in humans of 5 days, an extrinsic latent period (time from infection to infectiousness within the mosquito) of 10 days, and a mean infectious period in mosquitoes of 10 days. Under-reporting is assumed to be high (only 10% of cases reported) at the start of the outbreak and full reporting is assumed to be achieved in four weeks after the outbreak begins to grow. The estimated *R*_0_ for the Zika outbreak in San Andres was 1·41 (95% CI 1·15 to 1·74), and the *R*_0_ in Girardot was 4·61 (95% CI 4·11 to 5·16) (Table 2 and Figure 6). Odds ratios for gender and age effects were obtained from the likelihood model, indicating increased odds of transmission among females and adults aged 20 to 49 years old in both San Andres and Girardot (Table 2).

The estimation procedure was also applied to daily incidence data from a published outbreak in Salvador, Brazil, that occurred between February 15, 2015, and June 25, 2015; 14,835 cases were reported with an overall attack rate of 5·5 cases per 1,000 Salvador residents [7]. The estimated *R*_0_ of the Zika outbreak in Salvador, Brazil was 1·42 (95% CI 1·35 to 1·49). Sensitivity analyses are reported in the Supplementary Online Materials, including varying the incubation period in humans, the infectious period in humans, the infectious period in mosquitoes, the duration of under-reporting, and the level of under-reporting at the start of the outbreak.

## 4. Discussion

We report surveillance data on ZIKV outbreaks in two regions in Colombia between September 2015 and January 2016. The first region, San Andres, is a small, densely-populated island that is relatively isolated from continental Colombia. The second region, Girardot, is a typical moderately sized Colombian municipality. Both regions have endemic transmission of dengue and experienced recent outbreaks of CHIKV. We describe key epidemiological features of the Zika outbreaks and estimate the *R*_0_ from daily incidence data.

**Figure 6:**
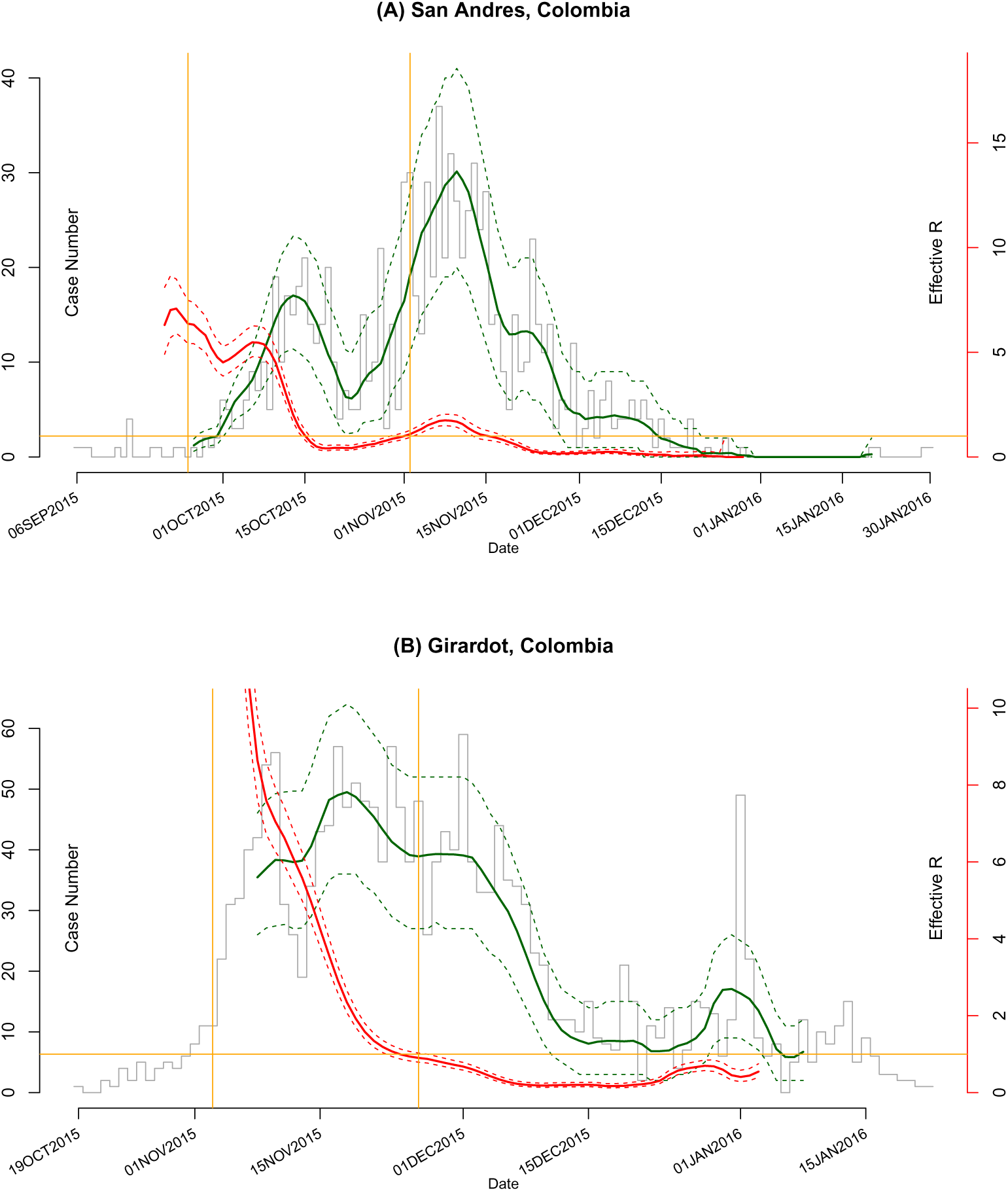
*(a)* Estimates of effective *R* (red) and model-fitted daily case numbers (green) for the outbreak of ZVD in San Andres, Colombia The reporting ratio is assumed to increase linearly from 10% on and before September 30, 2015, to 100% in 4 weeks. Dashed curves (both red and green) are conservative 95% CIs. Histogram in grey shows the epidemic curve. The horizontal orange line indicates the reference value of 1. The two vertical orange lines indicate the time interval used for the estimation of *R*_0_. *(b)* As (a) for Girardot, Colombia. The reporting ratio increases on October 19, 2015.

**Table 2:** Estimates of *R*_0_, gender-specific odds ratios for transmission, and age-specific odds ratios for transmission for ZVD in San Andres and Girardot, Colombia.

| Parameter | Level | San Andres |  | Girardot |  |
| --- | --- | --- | --- | --- | --- |
|  |  | Est. | 95% CI | Est. | 95% CI |
| $R_0$ | | 1.41 | (1.15, 1.74) | 4.61 | (4.11, 5.16) |
| $OR_{gender}$ | Male | - | - | - | - |
|  | Female | 1.71 | (1.50, 1.95) | 1.28 | (1.17, 1.40) |
| $OR_{age}$ | 20-49 | - | - | - | - |
|  | 0-19 | 0.86 | (0.74, 0.99) | 0.37 | (0.33, 0.42) |
| | $\geq 50$ | 0.74 | (0.63, 0.88) | 0.46 | (0.41, 0.52) |

The overall attack rates for ZVD as detected by local surveillance were 12·13 cases per 1,000 residents of San Andres and 18·43 cases per 1,000 residents of Girardot. These attack rates are similar to those reported from Yap Island (14·3 per 1,000) [3] but higher than those reported in Salvador, Brazil (5·5 per 1,000) [7]. In both areas, significantly higher attack rates are observed among women, especially those of child-bearing age. The Colombian government issued an epidemiological alert on December 2015 to actively search for pregnant women with Zika-like symptoms in areas with active transmission [27, 28]. This effort may partially explain the findings, though differences in gender-specific attack rates persist when only cases occurring prior to December are considered. Cases occurred in all age groups, but the most affected age group was 20 to 49 year of age, similar to previously published outbreaks in Yap Island, Federated States of Micronesia, and in Salvador, Brazil [3, 7]. As the population was fully susceptible to Zika transmission before the outbreaks, it is reasonable that all age groups would be affected.

Forty-eight pregnant women with ZVD were reported from San Andres and Girardot. These women are being followed according to national guidelines [27, 28] with no confirmed cases of microcephaly observed yet. Seventeen neurological syndromes, including GBS and ZIKV-associated meningoencephalitis, were identified, similar to reports from French Polynesia and Brazil [12, 29]. Laboratory-confirmation of these cases is challenging because neurological symptoms generally appear two weeks after acute symptoms [30] at which time ZIKV diagnosis by RT-PCR is not possible and serological tests are unreliable because of cross-reactivity with dengue [31, 32]. As ZIKV can be detected in urine longer than in blood [33], using urine samples to confirm ZIKV in GBS cases may be an alternative [34]. These challenges underscore the need for reliable diagnostic tests that can detect ZIKV after the viremic period.

The basic reproductive number (*R*_0_) was estimated in each area using daily incidence data. Our estimated *R*_0_ for the Zika outbreak in San Andres was 1·41 (95% CI 1·15 to 1·74), and the *R*_0_ for Girardot was 4·61 (95% CI 4·11 to 5·16). Applying the same methods with previously published data, we estimated that the *R*_0_ for Zika virus in Salvador, Brazil, was 1·42 (95% CI 1·35 to 1·49) [7]. We consider the estimate from San Andres to be the most reliable because it is a small, densely populated island, while Girardot has a higher risk of importation because the population fluctuates during weekends and holidays. The relative magnitudes of *R*_0_ are consistent with the higher dengue transmission observed in Girardot versus San Andres. Estimates of *R*_0_ in Zika are not widely available, though reports suggest an *R*_0_ of 4·3 to 5·8 in Yap Island and *R*_0_ of 1·8 to 2·0 in French Polynesia [35]. A recent manuscript considering the French Polynesian outbreak reported a range from 1·9 to 3·1 [36].

Relatively few cases were laboratory confirmed. The majority of cases were clinically confirmed, and the symptoms could be caused by other etiologies such as dengue. This report only includes symptomatic cases who attended a health care facility and were captured by the surveillance systems. ZIKV usually causes relatively mild illness lasting several days, and around 80% of infections are currently believed to be asymptomatic, so we are likely missing many mild or asymptomatic cases [10]. We also do not have a reliable estimate of under-reporting at these sites. Early under-reporting appeared to be especially apparent in the Girardot outbreak, and the sharp increase in cases observed may be due to increased public awareness of the disease. This phenomenon can result in an overestimate of *R*_0_.

Well-designed studies can provide valuable insight. Phylogenetic analyses of circulating ZIKV strains will be critical for understanding whether mutations in the viral genome are associated with an increased severity of disease, as manifested by microcephaly and GBS in this outbreak. Household studies can allow for more accurate estimation of transmission dynamics and enhance understanding of asymptomatic infection. Studies are required to understand the interactions between ZIKV, dengue, CHIKV, and other co-circulating arboviruses and their impact on disease. It is also necessary to increase surveillance of neurological syndromes associated with ZVD, like GBS and encephalitis.

The evidence for a causal relationship between ZIKV and microcephaly is strengthening [37, 38, 39]. Recent evidence from the French Polynesia outbreak suggests an estimated number of microcephaly cases associated with ZIKV is around one per 100 women infected in the first trimester[40]. Currently the Colombian Government is following a cohort of pregnant women that reported Zika-like symptoms anytime during their pregnancy. Those who are detected during the acute phase are being diagnosed with ZIKV RT-PCR. All women will be followed until the end of pregnancy, and the fetus will be evaluated during pregnancy and post-natally for twelve months. The prospective collection of data through this and other similar national cohorts will be essential for assessing causality, determining risk factors, and estimating rates of birth defects.

The results of this and other reports concludes that transmission of ZIKV may be widespread. Vector control has had limited success in controlling other arboviruses, such as dengue. A safe and efficacious vaccine, especially for women of child-bearing age, may be needed to reduce the disease burden.

## References

[1] J. E. Staples, E. J. Dziuban, M. Fischer, J. D. Cragan, Interim guidelines for the evaluation and testing of infants with possible congenital Zika virus infection — United States, 2016, MMWR Morb Mortal Wkly Rep 65 (2016) 63–67.

[2] A. D. Haddow, A. J. Schuh, C. Y. Yasuda, M. R. Kasper, V. Heang, R. Huy, H. Guzman, R. B. Tesh, S. C. Weaver, Genetic characterization of Zika virus strains: Geographic expansion of the Asian lineage, PLoS Negl Trop Dis 6 (2012) e1477.

[3] M. R. Duffy, T.-H. Chen, W. T. Hancock, A. M. Powers, J. L. Kool, R. S. Lanciotti, M. Pretrick, M. Marfel, S. Holzbauer, C. Dubray, L. Guillau-mot, A. Griggs, M. Bel, A. J. Lambert, J. Laven, O. Kosoy, A. Panella, B. J. Biggerstaff, M. Fischer, E. B. Hayes, Zika virus outbreak on Yap Island, Federated States of Micronesia, N Engl J Med 360 (2009) 2536–2543. PMID: 19516034.

[4] E. B. Hayes, Zika virus outside Africa 15 (2009) 1347.

[5] S. Ioos, H.-P. Mallet, I. L. Goffart, V. Gauthier, T. Cardoso, M. Herida, Current Zika virus epidemiology and recent epidemics, Medecine et Maladies Infectieuses 44 (2014) 302 – 307.

[6] J. Tognarelli, S. Ulloa, E. Villagra, J. Lagos, C. Aguayo, R. Fasce, B. Parra, J. Mora, N. Becerra, N. Lagos, L. Vera, B. Olivares, M. Vilches, J. Fernández, A report on the outbreak of Zika virus on Easter Island, South Pacific, 2014, Archives of Virology (2015) 1–4.

[7] C. W. Cardoso, I. A. Paploski, M. Kikuti, M. S. Rodrigues, M. M. Silva, G. S. Campos, S. I. Sardi, U. Kitron, M. G. Reis, G. S. Ribeiro, Outbreak of exanthematous illness associated with Zika, Chikungunya, and dengue viruses, Salvador, Brazil, Emerging Infectious Diseases 21 (2015) 2274–2276.

[8] Pan-American Health Organization, World Health Organization, Geographic distribution of confirmed cases of Zika virus (locally acquired) in countries and territories of the Americas, 2015-2016, 2016.

[9] European Centre for Disease Prevention and Control, Rapid risk assessment: Zika virus infection outbreak, French Polynesia, 2016.

[10] E. E. Petersen, J. E. Staples, D. Meaney-Delman, M. Fischer, S. R. Ellington, W. M. Callaghan, D. J. Jamieson., Interim guidelines for pregnant women during a Zika virus outbreak — United States, 2016, Morb Mortal Wkly Rep 65 (2016) 30–33.

[11] D. Musso, Zika virus transmission from French polynesia to Brazil, Emerging Infectious Diseases 21 (2015).

[12] V.-M. Cao-Lormeau, A. Blake, S. Mons, S. Lastére, C. Roche, J. Vanhomwegen, T. Dub, L. Baudouin, A. Teissier, P. Larre, A.-L. Vial, C. Decam, V. Choumet, S. K. Halstead, H. J. Willison, L. Musset, J.-C. Manuguerra, P. Despres, E. Fournier, H.-P. Mallet, D. Musso, A. Fontanet, J. Neil, F. Ghawché, Guillain-Barre syndrome outbreak associated with Zika virus infection in French Polynesia: a case-control study, The Lancet (2016).

[13] L. Schuler-Faccini, E. M. Ribeiro, I. M. Feitosa, D. D. Horovitz, D. P. Cavalcanti, A. Pessoa, M. J. R. Doriqui, J. I. Neri, J. M. de Pina Neto, H. Y. Wanderley, M. Cernach, A. S. El-Husny, M. V. Pone, C. L. Serao, M. T. V. Sanseverino, Possible association between Zika virus infection and microcephaly - Brazil, 2015, MMWR Morb Mortal Wkly Rep 65 (2016) 59–62.

[14] H. Lazear, E. Stringer, A. de Silva, The emerging Zika virus epidemic in the Americas: Research priorities, JAMA (2016).

[15] Instituto Nacional de Salud, Boletin epidemiologico semanal - semana epidemiologica 08 de 2016, 2016.

[16] Instituto Nacional de Salud, Vigilancia intensificada de síndromes neurológicos, (Guillain-Barre, polineuropatias ascendentes, entre otras afec-ciones neurologicas similares), en la fase epidémica de la infección por virus Zika en Colombia, 2015 – 2016, 2016.

[17] J. C. Padilla, D. P. Rojas, R. Saenz-Gomez, Dengue en Colombia: Epi-demiologia de la reemergencia a la hiperendemia, Guias de Impresion, first edition edition, 2012.

[18] Ministerio de Salud, Analisis de situacion de salud (ASIS) Archipielago de San Andres, 2011.

[19] D. Salas, Informe final del evento Chikungunya, Colombia 2014, 2014.

[20] Departamento Administrativo Nacional de Estadistica, Estimación y proyección de población Nacional, Departamental y Municipal total por área 1985–2020, 2016.

[21] T. García-Betancourt, D. R. Higuera-Mendieta, C. González-Uribe, S. Cortés, J. Quintero, Understanding water storage practices of urban residents of an endemic dengue area in Colombia: Perceptions, rationale and socio-demographic characteristics, PLoS ONE 10 (2015) 1–19.

[22] L. Alcala, J. Quintero, C. Gomez-Uribe, H. Brochero, Productividad de Aedes aegypti (l.) (Diptera: Culicidae) en viviendas y espacios públicos en una ciudad end éndemica para dengue en colombia, Biomedica 35 (2015).

[23] Ministerio de Salud, Instituto Nacional de Salud, Circular conjunta externa no. 061 de 2015, 2015.

[24] Instituto Nacional de Salud, Report form Zika illness-Colombia, 2015.

[25] R Core Team, R: A Language and Environment for Statistical Computing, R Foundation for Statistical Computing, Vienna, Austria, 2015.

[26] Y. Yang, J. D. Sugimoto, M. E. Halloran, N. E. Basta, D. L. Chao, L. Matrajt, G. Potter, E. Kenah, I. M. Longini, The transmissibility and control of pandemic Influenza A (H1N1)virus, Science (2009).

[27] Ministerio de Salud, Instituto Nacional de Salud, Circular externa 004 de 2016, 2016.

[28] Instituto Nacional de Salud, Circular 007 de 2016, 2016.

[29] World Health Organization, Guillain-Barrú syndrome - Brazil, 2016.

[30] N. Yuki, H.-P. Hartung, Guillain-Barrú syndrome, New England Journal of Medicine 366 (2012) 2294–2304. PMID: 22694000.

[31] G. S. Campos, A. C. Bandeira, S. I. Sardi, Zika virus outbreak, Bahia, Brazil, Emerging Infectious Diseases 21 (2015) 1885–1886.

[32] W. E. Villamil-Gómez, O. González-Camargo, J. Rodriguez-Ayubi, D. Zapata-Serpa, A. J. Rodriguez-Morales, Dengue, chikungunya and Zika co-infection in a patient from Colombia, Journal of Infection and Public Health (2016).

[33] A.-C. Gourinat, O. O’Connor, E. Calvez, C. Goarant, M. Dupont-Rouzeyrol, Detection of Zika virus in urine, Emerging Infectious Diseases 21 (2015).

[34] B. Rozé, F. Najioullah, J. Ferge, K. Apetse, Y. Brouste, R. Cesaire, C. Fagour, L. Fagour, P. Hochedez, S. Jeannin, J. Joux, H. Mehdaoui, R. Valentino, A. Signate, A. Cabie, Zika virus detection in urine from patients with Guillain-Barré syndrome on Martinique, January 2016, Euro Surveillance 21 (2016).

[35] H. Nishiura, R. Kinoshita, K. Mizumoto, Y. Yasuda, K. Nah, Transmission potential of zika virus infection in the south pacific, International Journal of Infectious Diseases (2016) -.

[36] A. J. Kucharski, S. Funk, R. M. M. Eggo, H.-P. Mallet, J. Edmunds, E. J. Nilles, Transmission dynamics of Zika virus in island populations: a modelling analysis of the 2013-14 French Polynesia outbreak, bioRxiv (2016).

[37] J. Mlakar, M. Korva, N. Tul, M. Popović, M. Poljšak-Prijatelj, J. Mraz, M. Kolenc, K. Resman Rus, T. Vesnaver Vipotnik, V. Fabjan Vodušek, A. Vizjak, J. Pižem, M. Petrovec, T. Avšiĉ Županc, Zika virus associated with microcephaly, New England Journal of Medicine 374 (2016) 951–958. PMID: 26862926.

[38] World Health Organization, Zika situation report, 2016.

[39] P. Brasil, J. P. Pereira, Jr., C. Raja Gabaglia, L. Damasceno, M. Wakimoto, R. M. Ribeiro Nogueira, P. Carvalho de Sequeira, A. Machado Siqueira, L. M. Abreu de Carvalho, D. Cotrim da Cunha, G. A. Calvet, E. S. Neves, M. E. Moreira, A. E. Rodrigues Baião, P. R. Nassar de Carvalho, C. Janzen, S. G. Valderramos, J. D. Cherry, A. M. Bispo de Filippis, K. Nielsen-Saines, Zika virus infection in pregnant women in Rio de Janeiro — preliminary report, New England Journal of Medicine 0 (2016) null. PMID: 26943629.

[40] S. Cauchemez, B. P. Besnard, Marianne AND, T. Dub, P. Guillemette-Artur, D. Eyrolle-Guignot, et al., Association between zika virus and microcephaly in french polynesia, 2013-15: a retrospective study, The Lancet 0 (2016) DOI: http://dx.doi.org/10.1016/S0140-6736(16)00651-6.

